# Resistance genes are distinct in protein-protein interaction networks according to drug class and gene mobility

**DOI:** 10.1101/2024.02.05.578986

**Authors:** Nazifa Ahmed Moumi, Connor L. Brown, Shafayat Ahmed, Peter J. Vikesland, Amy Pruden, Liqing Zhang

## Abstract

With growing calls for increased surveillance of antibiotic resistance as an escalating global health threat, improved bioinformatic tools are needed for tracking antibiotic resistance genes (ARGs) across One Health domains. Most studies to date profile ARGs using sequence homology, but such approaches provide limited information about the broader context or function of the ARG in bacterial genomes. Here we introduce a new pipeline for identifying ARGs in genomic data that employs machine learning analysis of Protein-Protein Interaction Networks (PPINs) as a means to improve predictions of ARGs while also providing vital information about the context, such as gene mobility. A random forest model was trained to effectively differentiate between ARGs and nonARGs and was validated using the PPINs of ESKAPE pathogens (*Enterococcus faecium, Staphylococcus aureus, Klebsiella pneumoniae, Acinetobacter baumannii, Pseudomonas aeruginosa*, and *Enterobacter cloacae*), which represent urgent threats to human health because they tend to be multi-antibiotic resistant. The pipeline exhibited robustness in discriminating ARGs from nonARGs, achieving an average area under the precision-recall curve of 88%. We further identified that the neighbors of ARGs, i.e., genes connected to ARGs by only one edge, were disproportionately associated with mobile genetic elements, which is consistent with the understanding that ARGs tend to be mobile compared to randomly sampled genes in the PPINs. This pipeline showcases the utility of PPINs in discerning distinctive characteristics of ARGs within a broader genomic context and in differentiating ARGs from nonARGs through network-based attributes and interaction patterns. The code for running the pipeline is publicly available at https://github.com/NazifaMoumi/PPI-ARG-ESKAPE

## Introduction

The increasing prevalence of antibiotic-resistant infections poses a significant health threat(1). Previously treatable diseases are now becoming untreatable due to the evolution and spread of resistant bacterial strains(2). According to the Centers for Disease Control and Prevention (CDC), more than 2.8 million people are infected with antibiotic-resistant pathogens annually, resulting in approximately 35,000 deaths(3; 4; 5). Of particular concern is the evolution and spread of novel resistance phenotypes, which results in an ever-evolving challenge to identify effective antibiotics in the clinic and maintain their efficacy for years to come. Novel resistance phenotypes can result from the horizontal transfer of antibiotic resistance genes (ARGs) to new species or strains or through previously unknown ARGs emerging in the genome(6; 7).

The advent of next-generation DNA sequencing over the past decade represents a promising approach to support One Health surveillance of antibiotic resistance, i.e., across humans, animals, plants/crops, and the environment. Whole genome sequencing can be applied to profile the dominant genes across the microbial community inhabiting an environment of interest (e.g., sewage, manure, soil, food). These sequences can then be compared against publicly available databases to profile ARGs and thus profile and compare genotypic resistance patterns. However, the incompleteness of public databases is an inherent limitation of this approach, particularly if there is interest in monitoring previously unidentified ARGs(8). False positives are also possible, due to local sequence similarity(9). Also, simple read-matching homology-based profiling of this nature ignores the context of putative ARGs and other genes of importance in potentiating and mobilizing antibiotic resistance.

DNA sequences can be assembled into longer contiguous sequences to provide more complete information about the ARG of interest and thus increase confidence in its annotation and functional assignment when compared to public databases(10). Machine learning approaches, especially deep learning, can also help to improve the prediction of ARGs, including novel ARGs, relative to simple sequence homology-based comparisons(11; 12; 13; 14). However, such approaches still fail to tap into broader information available in the genome to precisely predict ARG function(15). These approaches struggle to effectively encompass complex interactions between various ARGs as well as genes of divergent types(16). They also provide little information about other genes involved in the mobility of the ARG or expression of its phenotype(17; 18). Focusing analysis on proteins, instead of nucleotide sequences, could present advantages in this regard because it is the proteins encoded by ARGs that perform the ultimate function of conferring resistance to antibiotics, e.g., by target modification, enzymatic degradation of the antibiotic, or pumping the antibiotic out of the cell. In particular, analysis of protein-protein interaction networks (PPINs) could help to address this need, by enhancing understanding of the proteins involved in resistance, and their interactions within an organism(19).

The overall objective of this study was to develop and validate a PPIN-based pipeline for characterizing ARGs in whole genome data. We hypothesized that ARGs would exhibit distinct patterns in network topology relative to nonARGs in the PPIN, which can then be recognized by machine-learning algorithms to predict their resistance mechanisms. Of particular interest was to assess whether ARGs are likely to be mobile, based on the strength of their networks with mobile genetic elements (MGEs). To validate the pipeline, we analyzed whole genome sequences of representative “ESKAPE” pathogens (*Enterococcus faecium, Staphylococcus aureus, Klebsiella pneumoniae, Acinetobacter baumannii, Pseudomonas aeruginosa*, and *Enterobacter cloacae*), which represent an urgent clinical threat because of their tendency to be multi-antibiotic resistant due to carriage of multiple ARGs on MGEs(20; 21; 22). The findings of this study highlight the potential of PPIN-based analysis as a new and accurate means of classifying ARGs, providing much more comprehensive characterization than typical nucleotide sequence read-matching approaches. The approach here also overcomes the limitations of publicly available databases, enabling the discovery of previously unknown ARGs.

## Materials and methods

The experimental steps are depicted in Fig. 1, providing a visual overview of the process.

**Fig. 1.**
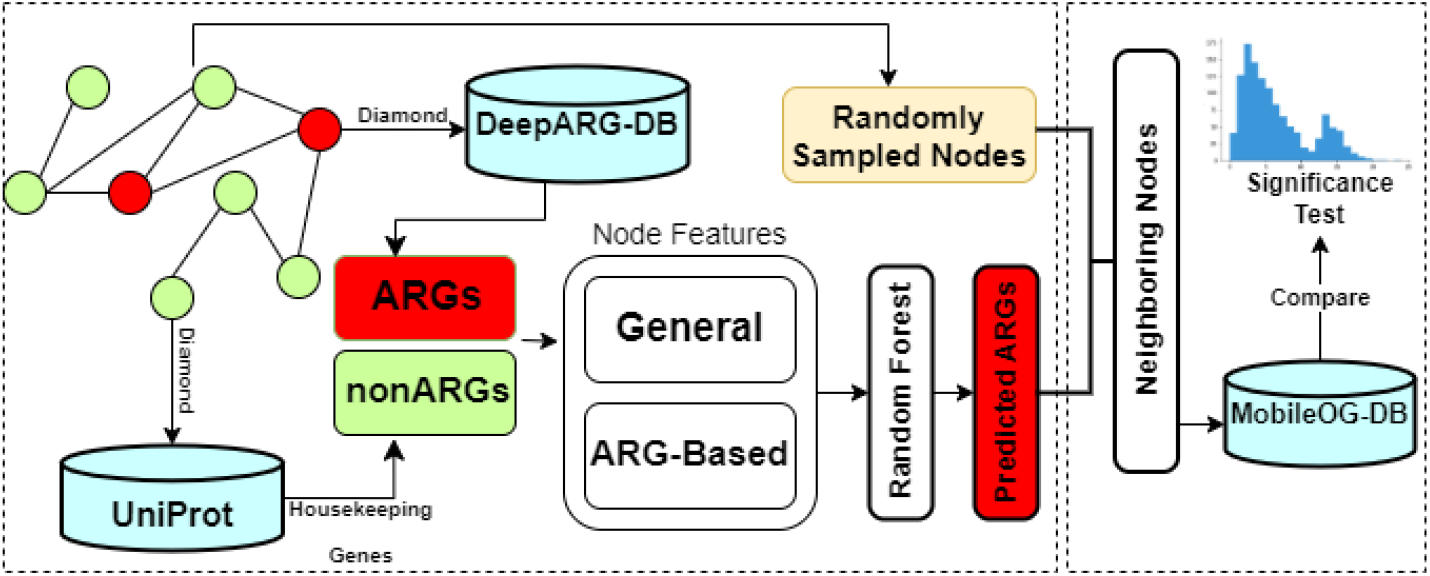
Overview of the pipeline. A random forest classifier is used to predict the ARGs in the Protein-Protein Interaction Networks (PPIN). NonARGs are randomly sampled and annotated using MobileOG-DB for comparing their mobility relative to ARGs

### Data collection

The PPINs for ESKAPE pathogens were collected from the String DB(23). The interaction type used for the analysis is physical and every interaction has a confidence score associated with it. If proteins exhibit indications of co-occurrence within a protein complex, physical interaction scores are calculated based on selected evidence channels. These scores are then combined to create an aggregated physical interaction score. The resulting physical interaction score represents the probability of two proteins being present together in a gold-standard set of protein complexes. We opted for the physical network instead of the functional interaction network due to the higher confidence scores associated with physical interactions.

Proteins that are likely to encode antibiotic resistance were selected from the set of proteins in the PPINs by using DIAMOND alignment with the reference ARG database from DeepARG(25; 11). There are 14,872 ARGs in the latest DeepARG-DB. After aligning with an identity cutoff of 70%, varying numbers of ARGs were detected in the PPINs of the pathogens (Table 1). To derive the negative control set of nonARGs, we reasoned that housekeeping genes are less likely to play a specific role in drug resistance, especially those that encode the enzymes that are necessary for basic metabolic processes(26), and therefore selected genes from 184 different Gene Ontology (Biological Process) terms that cover some of the well-established characteristics of housekeeping genes, such as different metabolic processes, excision repair of nucleotides, aerobic respiration chain, etc(27; 28) (Supplementary Table 1). By listing the proteins from the selected pathogens from UniProt that contain at least one of these GO terms in its functionality, the designated housekeeping genes are found for these microorganisms, which, in the following process, are used as the negative set.

**Table 1.** Network Information (total number of proteins/nodes, interactions/edges, ARGs, neighboring nodes to the ARGs in the PPINs) for ESKAPE pathogens from STRING-DB.

|  | *E. faecium* | *S. aureus* | *A. baumannii* | *K. pneumoniae* | *P. aeruginosa* | *E. cloacae* |
| --- | --- | --- | --- | --- | --- | --- |
| #Proteins (Nodes) | 1,867 | 1,834 | 2,229 | 3,688 | 3,897 | 3,384 |
| #Interactions (Edges) | 26,874 | 21,669 | 26,171 | 35,520 | 63,007 | 35,778 |
| #ARGs | 182 | 181 | 216 | 468 | 447 | 388 |
| #Neighbours to the ARG nodes | 857 | 797 | 825 | 1,177 | 1,504 | 1,264 |

### Feature selection

The PPIN can be represented as an undirected graph denoted by G(V, E), where V represents the set of protein vertices and E represents the set of edges. In this graph, proteins are connected by an edge if they interact with each other. To discern between ARGs and nonARGs in the PPIN, we selected ten network topology-based features calculated using NetworkAnalyzer in Cytoscape and five ARG node-based features(29) (Table 2,3 To further refine our feature selection by eliminating redundancy, we computed the pairwise correlations between the features. We have excluded any features that exhibit a correlation exceeding 0.95 with another feature (Supplementary Fig. 1).

**Table 2.** Topological Features Calculated by NetworkAnalyzer.

| Feature | Description |
| --- | --- |
| AverageShortestPathLength | Expected distance between two connected nodes |
| BetweennessCentrality | Control that this node exerts over the interactions of other nodes in the network |
| ClosenessCentrality | How fast information spreads from a given node to other reachable nodes in the network |
| ClusteringCoefficient | A measure of the degree to which nodes in a graph tend to cluster together |
| Degree | The number of edges linked to a node |
| Eccentricity | The maximum non-infinite length of the shortest path between a node and another in the network |
| NeighborhoodConnectivity | Average connectivity of all neighbors of a node |
| Radiality | A node centrality index |
| Stress | Counts the number of shortest paths passing through a node |
| TopologicalCoefficient | Quantify the extent to which a protein in the network shares interaction partners with other proteins |

**Table 3.**
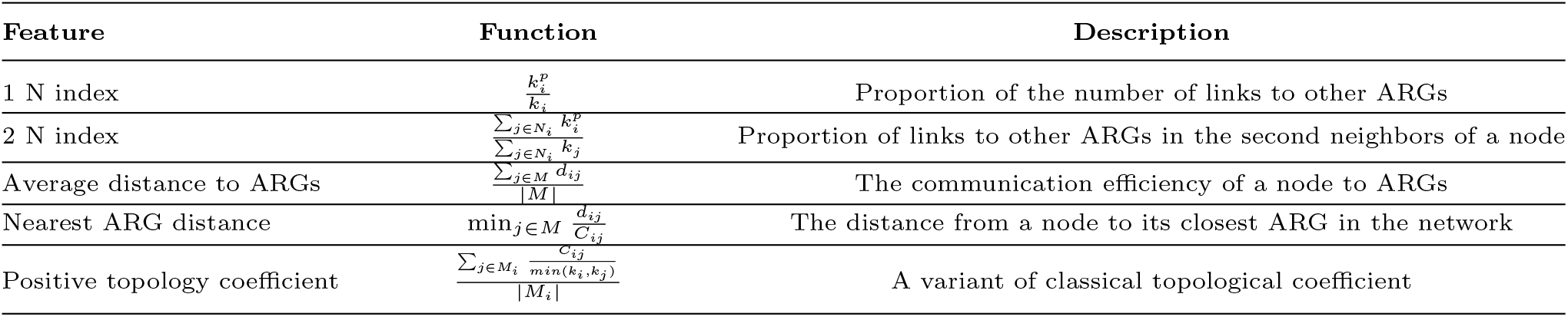
ARG node-based features.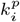 denotes the number of edges lined to an ARG; *k*_*i*_ denotes the degree of the node; *N*_*i*_ denotes a node-set consisting of all the neighbors of node *i*; *M* denotes a node set consisting of all the ARGs; *d*_*ij*_ denotes the shortest path between two nodes; *C*_*ij*_ denotes the number of nodes that are connected to both nodes *i* and *j*; *M*_*i*_ denotes a node set consisting of nodes that share neighbors with node *i*.

| Feature | Function | Description |
| --- | --- | --- |
| 1 N index | $\frac{k_i^p}{k_i}$ | Proportion of the number of links to other ARGs |
| 2 N index | $\frac{\sum_{j \in N_i} k_j^p}{\sum_{j \in N_i} k_j}$ | Proportion of links to other ARGs in the second neighbors of a node |
| Average distance to ARGs | $\frac{\sum_{j \in M} d_{ij}}{\vert M\vert }$ | The communication efficiency of a node to ARGs |
| Nearest ARG distance | $\min_{j \in M} \frac{d_{ij}}{C_{ij}}$ | The distance from a node to its closest ARG in the network |
| Positive topology coefficient | $\frac{\sum_{j \in M_i} \frac{C_{ij}}{\min(k_i, k_j)}}{\vert M_i\vert }$ | A variant of classical topological coefficient |

### Random forest classifier

The constructed networks for ESKAPE pathogens had insufficient data, thereby limiting the use of deep learning frameworks that require extensive training data to produce satisfactory results. As such, we employed a random forest classifier with the number of trees determined empirically. The Random Forest model includes a built-in feature selection mechanism calculating the decrease in node impurity weighted by the probability of reaching that node, enabling us to identify the most relevant features crucial for accurately distinguishing ARGs from nonARGs. Moreover, it is well-suited for analyzing high-dimensional biological data due to its robustness to noise and outliers(30). By leveraging ensemble learning, Random Forest reduces overfitting and improves generalization performance, which is especially valuable when working with limited sample sizes compared to the number of features.

To enhance the prediction performance of our model, we implemented a sampling technique (Fig. 2). In this approach, we considered M as the number of instances in the minority class (ARG nodes) and N as the number of instances in the majority class (nonARG nodes) within the training dataset, with M significantly smaller than N. During each iteration, we randomly sampled M instances from the majority class. Subsequently, we combined these M instances with all instances from the minority class to train one Random Forest model. This sampling process was repeated k times to train k separate models and eventually, all of the predictions were ensembled together. By employing this sampling method, we ensured that each instance in the majority class was selected and trained alongside an equal number of instances from the minority class, mitigating the risk of overfitting. As such, we achieved a balanced 1:1 ratio between ARGs and nonARGs by creating several undersampled subsets within the nonARGs class. Each of these subsets served as the training data for separate Random Forest models. Later, we amalgamated the predictions from these individual models through a majority voting approach.

**Fig. 2.**
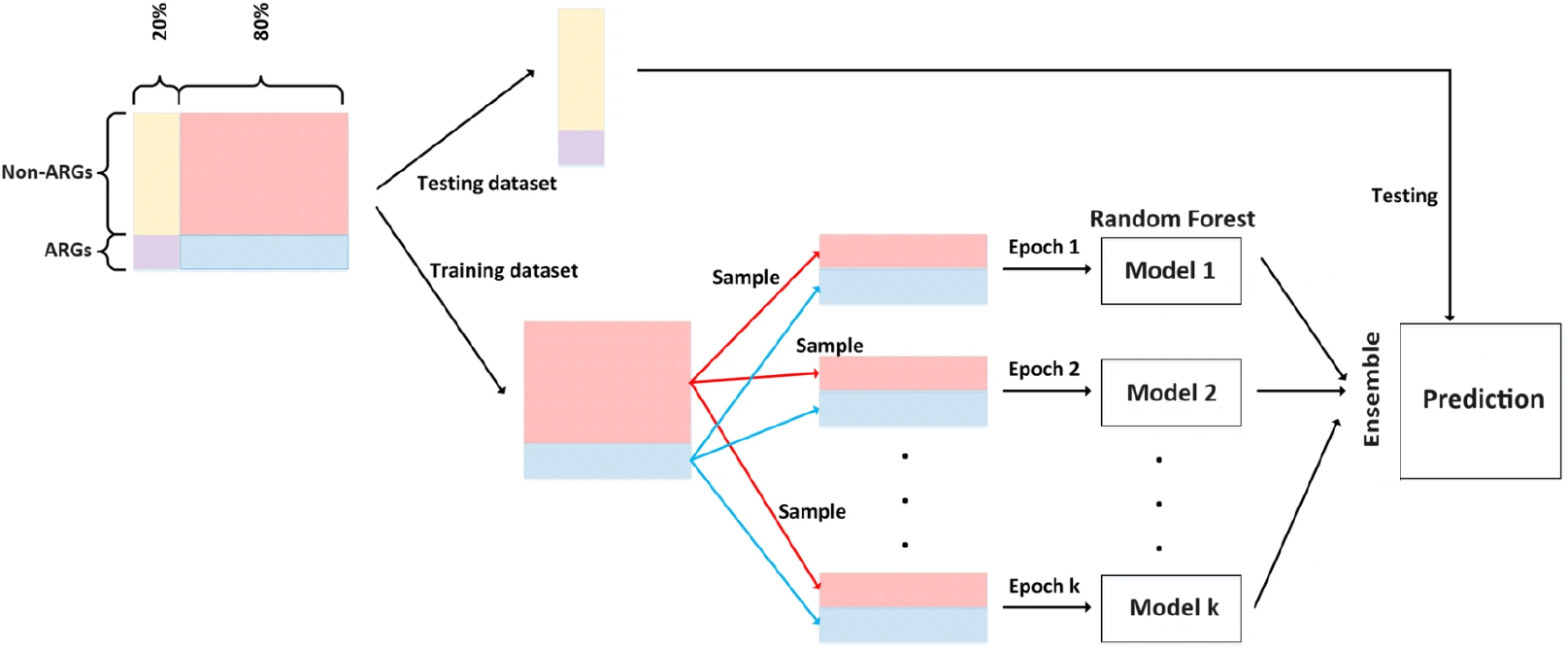
Illustration of the random sampling approach applied to both ARG and non-ARG sets to achieve a balanced 1:1 ratio. Following this, the prediction results obtained from these samples were aggregated for the final prediction score

We split our dataset into an 80-20 ratio for training and testing and ensured that the ARGs on the testing set were excluded while calculating the features. This exclusion minimized potential bias in the feature values calculated from the network specifically for the ARG-based features. We assessed the performance of our random forest classifier by training six models on six different pathogens. We varied the number of trees in each model from 10 to 400 and evaluated the precision and recall metrics to select the optimal number of trees to be used.

To rigorously evaluate the statistical significance of our models in accurately identifying antibiotic resistance genes (ARGs), we established a null hypothesis framework. This involved analyzing the distribution of evaluation metric scores derived from ten randomly selected positive sets, which represented ARGs within the network. For each organism under study, we systematically carried out a random sampling procedure to generate the same number of proteins as the actual ARGs identified specifically for that organism. This random sampling process was repeated ten times, yielding ten distinct sets of randomly selected positive proteins for each organism.

Subsequently, we trained separate Random Forest models by treating these randomly selected sets as the ARGs or positive sets, while the remaining proteins were considered nonARGs or negative sets. The sampling strategy was replicated for this analysis to ensure a 1:1 ratio between the positive and the negative sets. The prediction results obtained from these models were then aggregated using the average method. This approach enabled us to construct a robust statistical framework for assessing the performance of our models by comparing their results to the average evaluation scores derived from these randomized positive sets.

### Evaluation metrics

To obtain a reliable estimate of our model’s performance, we conducted a 5-fold cross-validation. The performance of our model was evaluated using the following metrics:

- **Accuracy** is the ratio of the number of correct predictions to the total number of predictions.
- **Precision** is the ratio of the number of correctly predicted ARGs to the number of predicted ARGs.
- **Recall** is the ratio of the number of correctly predicted ARGs to the total number of ARGs.
- **F1 Score** is the harmonic mean of precision and recall:

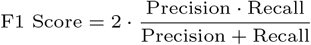

Additionally, we calculated the AUPRC (Area Under the Precision-Recall Curve) values for each of the six models as it is particularly useful when dealing with imbalanced data.

### Mobility analysis of nodes in PPINs

To gain insight into potential drivers of the dissemination of ARGs, we inspected the mobility of the ARGs as well as their first-order neighbors in the PPINs. We used mobileOG-DB as the reference database to label the proteins from the PPIN into two of the major categories from mobileOG, namely, Transfer and Excision/Integration(31). Proteins mediating the transfer mechanism or proteins associated with site-specific recombination mechanisms were annotated using this database.

We examined whether ARGs are more mobile compared to the other nodes by comparing the number of ARGs to linkages with MGE hallmark genes compared to randomly sampled genes. In addition, we aimed to examine the comparative mobility of genes resistant to different drug classes.

#### Determining the putative mobility of the ARGs in PPIN

To investigate the mobility of ARGs within the PPIN, we conducted an alignment of the proteins associated with the ARGs from the network against the mobileOG-DB using DIAMOND. This alignment aimed to identify potential mobile ARGs that possess either of two pertinent tags: “Transfer” or “Excision/Integration,” with a minimum identity threshold of 70%.

Subsequently, we proceeded to assess whether there was a statistically significant difference in the frequency of tags among the neighbors of ARGs, which refers to proteins directly interacting with ARGs, in comparison to other randomly sampled proteins. To assess the relative mobility of ARGs, we employed a sampling approach. We randomly selected an equal number of proteins matching the number of ARGs from the entire protein set for each organism. From the combined neighboring nodes of these randomly selected proteins, we again randomly sampled an equal number of proteins to match the number of neighbors to the ARGs for a specific organism. In cases where an insufficient number of neighbors were available, we repeated some of the proteins in the sampled set to maintain the desired sample size. This particular sampling technique was employed to ensure that the network structure remained consistent between the actual neighboring proteins of ARGs and the proteins chosen randomly. This sampling process was repeated 100 times. For each sample, we determined the mobility tags using DIAMOND alignment with mobileOG-DB (identity cutoff 70%). We then compared the distributions for the neighbors and non-neighbors and conducted a one-tailed t-test to evaluate the significance.

Because of the association of genes resistant to specific drug classes with MGEs, we hypothesized that the first-order neighbors of different families of ARGs would be disproportionately associated with MGE-hallmark genes. To further investigate the relative mobility of genes resistant to different drug classes, we applied the same sampling strategy for each drug class. We combined the protein sets from the six pathogens and identified genes resistant to a specific drug class of interest. We then extracted the neighborhoods of these genes and repeated the sampling process like before to obtain 100 random samples of genes, matching the number of neighbors for the group of genes resistant to that specific drug class. Significance tests were performed to compare the relative mobility with the randomly picked protein sets, as well as across different groups of genes resistant to different drug classes. This allowed us to assess the differences in mobility patterns among various drug classes and evaluate their significance.

## Results

### Overall network topology and model accuracy

The results of our analysis revealed an average Area Under the Precision-Recall Curve (AUPRC) of 0.8816 across all the ESKAPE pathogens, indicating relatively good overall performance (Fig. 3). However, the performance was better for some species rather than others. the Model trained on the network from *S. aureus* has the lowest AUPRC 0.813, likely due to the observation that its PPIN has the least number of proteins and also the least number of ARGs. By contrast, the model trained on the network from *K. pneumoniae* demonstrated the highest performance across all evaluation metrics (AUPRC: 0.914). Comparatively, *K. pneumoniae*’s PPIN has one of the largest number of nodes as well as the number of ARGs, and the increased network complexity may provide more meaningful patterns and information for accurate prediction.

**Fig. 3.**
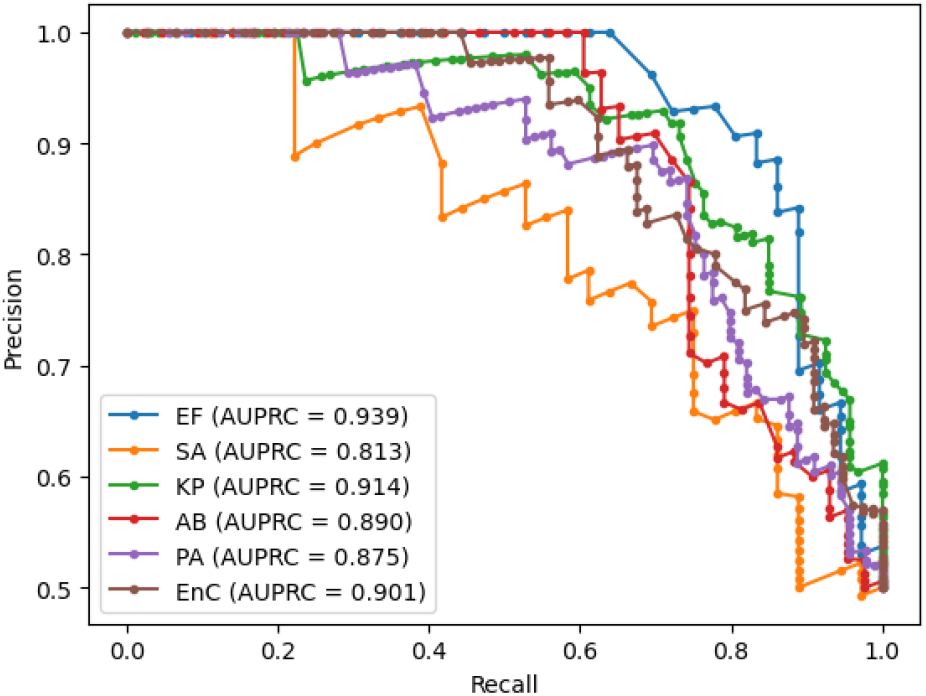
Area Under the Precision-Recall Curve (AUPRC) plot of the six ESKAPE pathogens(EF = *Enterococcus faecium*, SA = *Staphylococcus aureus*, KP = *Klebsiella pneumoniae*, AB = *Acinetobacter baumannii*, PA = *Pseudomonas aeruginosa*, EnC = *Enterobacter cloacae*).

In order to assess the statistical significance of our models’ performances, we implemented a null hypothesis by analyzing the distribution of evaluation metrics’ scores obtained from ten randomly chosen positive sets for each organism of interest, which were considered ARGs in the network. Across different organisms, we consistently observed lower values for accuracy, recall, precision, and f1-score, averaging around 45% (Fig. 4), for identifying the randomly sampled positive nodes. Importantly, when evaluating the model’s performance using the actual set of ARGs specific to each organism, we observed a substantial improvement in the evaluation metrics, nearly doubling their values.

**Fig. 4.**
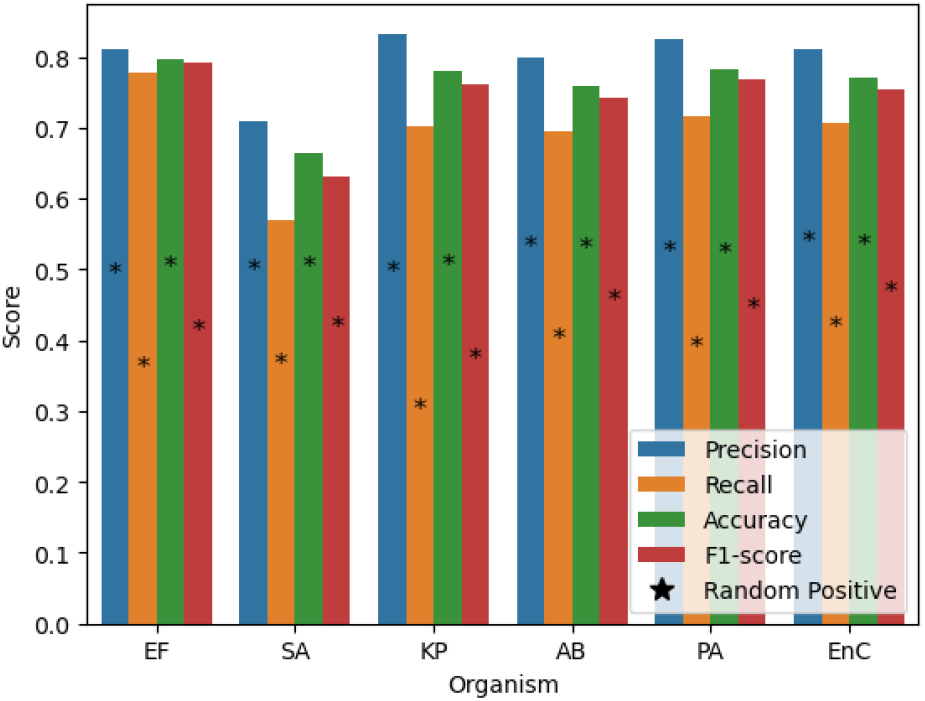
Performance evaluation of the ARG prediction model by four different metrics (precision, recall, accuracy, and F1 score) across the six pathogens; * marks indicate the evaluation scores obtained by considering a randomly picked set of proteins as the positive proteins.

Principal Component Analysis (PCA) was performed for all of the proteins across all of the ESKAPE pathogens by merging the topological feature sets in order to explore whether ARGs exhibit a tendency to cluster together based on network-based features. In the PCA plot (Supplementary Fig. 2), we observed a discernible separation between ARGs and nonARGs. The orange dots representing ARGs appeared predominantly in the top region of the plot, while the nonARGs were primarily located in the lower region. Furthermore, we performed single linkage hierarchical clustering to investigate the discernibility of different drug classes based on topological features of the PPIN. The clustering was conducted using the Mahalanobis distance between the means of the 10 PCs from the PCA for the drug classes. We focused on the top 20 most frequent drug classes identified across the six pathogens. The resulting dendrogram provided insights into the similarity and dissimilarity between the drug classes, visually depicting their hierarchical relationships (Supplementary Fig. 3).

In our multi-drug class classification, we observed that certain drugs such as macrolide, fluoroquinolone, multidrug, and tetracycline are highly prevalent across the ESKAPE pathogens (Supplementary Fig. 4). Notably, our model was able to classify these frequently occurring drugs, as evidenced by the darker shades of color along the diagonal in the heatmap of the confusion matrix (Fig. 5). In particular, glycopeptide, peptide, and macrolide drug classes were consistently present among the top ten correctly classified drug classes in all six pathogen-specific models. The majority of genes belonging to these drug classes were accurately identified by our models. However, it’s important to note that the performance of our model varied among different organisms. For instance, for the network from *K. pneumoniae*, the classification performance was generally better compared to networks from *S. aureus* and *A. baumannii*, which have relatively fewer proteins.

**Fig. 5.**
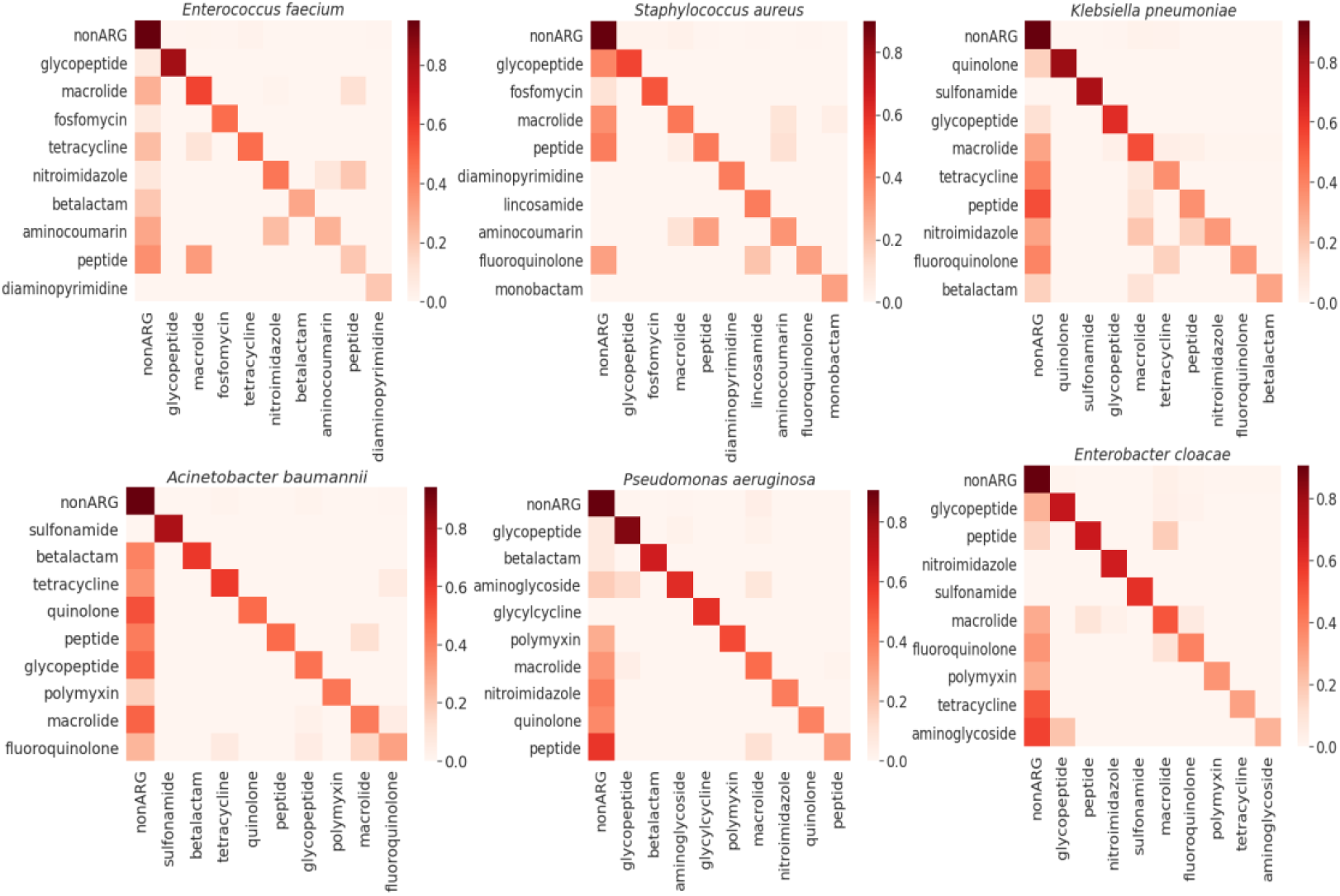
Normalized Confusion Matrix illustrating the performance of the PPIN-based model in classifying genes resistant to multiple drug classes based on the PPINs of the ESKAPE pathogens (The matrix displays true drug classes on the y-axis and predicted drug classes on the x-axis). The pathogens are listed from left to right representing *Enterococcus faecium, Staphylococcus aureus, Klebsiella pneumoniae, Acinetobacter baumannii, Pseudomonas aeruginosa*, and *Enterobacter cloacae*, respectively. The confusion matrix offers a visual depiction of the classification performance, showing the balance between accurate and erroneous predictions within the top 10 correctly predicted drug classes for each pathogen.

We also observed that some genes from less common drug classes were misclassified as nonARGs. This can be attributed to the limited number of representative genes from these classes in our training set. As a result, our model may not have learned sufficient interaction patterns to accurately classify these less prevalent drug classes.

The analysis of feature importance in predicting ARG nodes provides valuable insights into the interaction patterns of ARGs in the PPIN (Fig. 6(a)). Among the top five most informative features contributing to the accuracy of the model, two of them are centrality-based features. Stress centrality and betweenness centrality play a crucial role in facilitating the flow of information across the PPIN(32; 33). They determine the essentiality of a particular protein in carrying out an associated function by connecting bridges within the network. In the PPIN, many proteins’ shortest pathways pass through those with high centrality measures. They act as mediators between other proteins, catalyzing reactions or acting as intermediate substances. Alterations in these proteins (mutation) can affect the interactions between other proteins whose shortest pathways rely on them. By incorporating feature values from all ESKAPE pathogens, we can see that ARGs exhibit higher values for stress, closeness centrality, clustering coefficient, and neighborhood connectivity compared to nonARGs (Fig. 6(b)). A node with high stress in a PPIN is likely to be a central player in mediating interactions and communication within the network. These nodes are distinguished by a notably higher incidence of shortest paths traversing through them reflecting their importance in coordinating various biological functions.

**Fig. 6.**
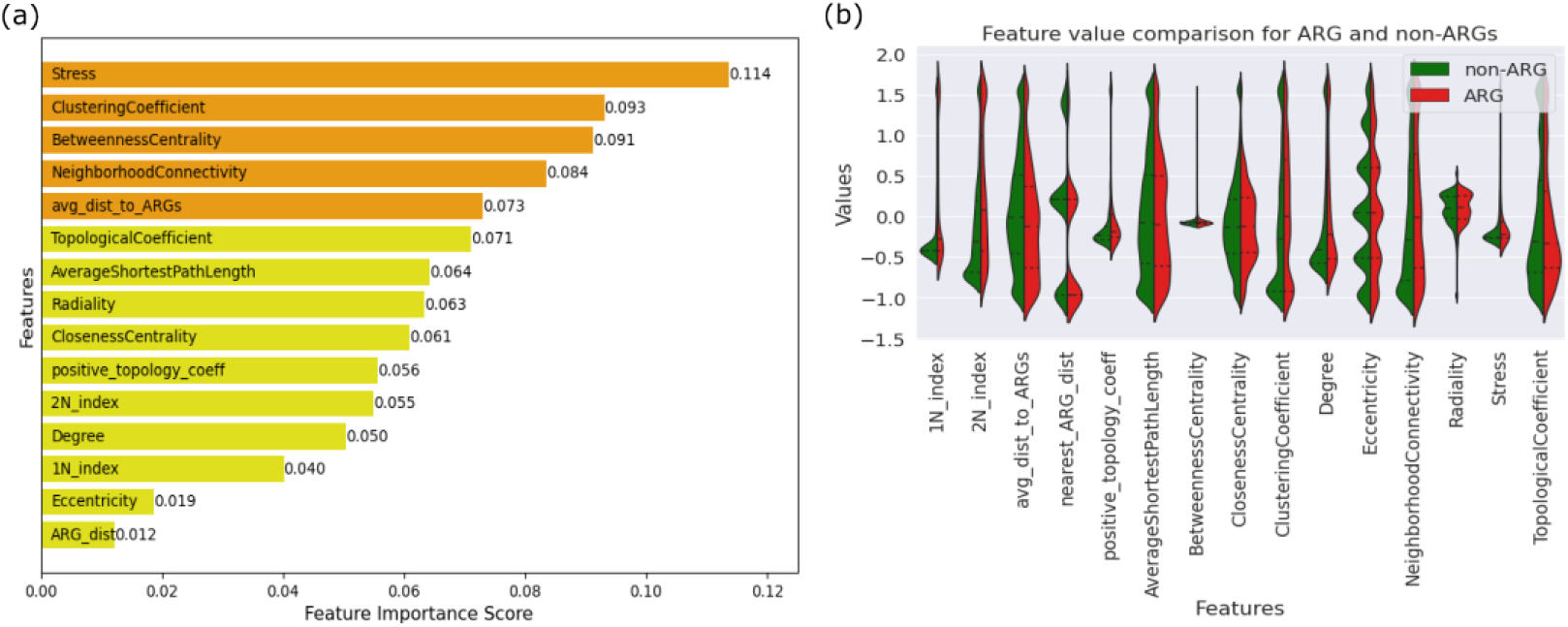
(a) Average feature importance to the model’s performance across all six pathogens. (b) The distribution of feature values for both ARGs and nonARGs

We can also observe an ARG-based feature: the average distance to ARG in the top five features. However, the values for this feature suggest that resistance genes typically do not exhibit a pattern of close proximity to other resistance genes within the PPIN. The positive topological coefficient, a variant of the general topological coefficient metric, measures the frequency of a node sharing common neighbors with ARG nodes. The 1N index and the 2N index calculate the ratio of ARGs to all neighboring proteins and to all the second neighbors respectively for a specific node; while the average ARG distance represents the mean length of the shortest paths between the query protein and ARGs. The relative importance of these ARG-based features suggests that protein interactions with ARGs and their shared neighborhoods are significant indicators of resistance in PPINs.

### Integration/excision and conjugation proteins in the PPINs show a significant association with ARGs

We performed mobility analysis of the 17 prevalent drug classes, including multidrug, tetracycline, fluoroquinolone, macrolide, glycopeptide, peptide, macrolide, aminoglycoside, monobactam, aminocoumarin, cephalosporin, fosfomycin, nucleoside, beta-lactam, polymixin, quinolone, phenicol, sulfonamide which are known to be associated with MGEs in ESKAPE pathogens(20). The plots indicate a significant and consistent association between ARGs from these prevalent drug classes and transfer/excision elements in the PPINs (Fig. 7). While drug classes including aminoglycoside, aminocoumarin, cephalosporin, monobactam, and multidrug showed a significant association with MGEs, drug classes such as sulfonamide, fosfomycin, nucleoside, phenicol, tetracycline, and glycylcycline exhibited a significantly lower association. Drug classes such as bicyclomycin, bacitracin, bleomycin, chloramphenicol, elfamycin, fosmidomycin, kasugamycin, lincosamide, mupirocin, and oxazolidinone were not included in the analysis due to the limited number of genes in the classes.

**Fig. 7.**
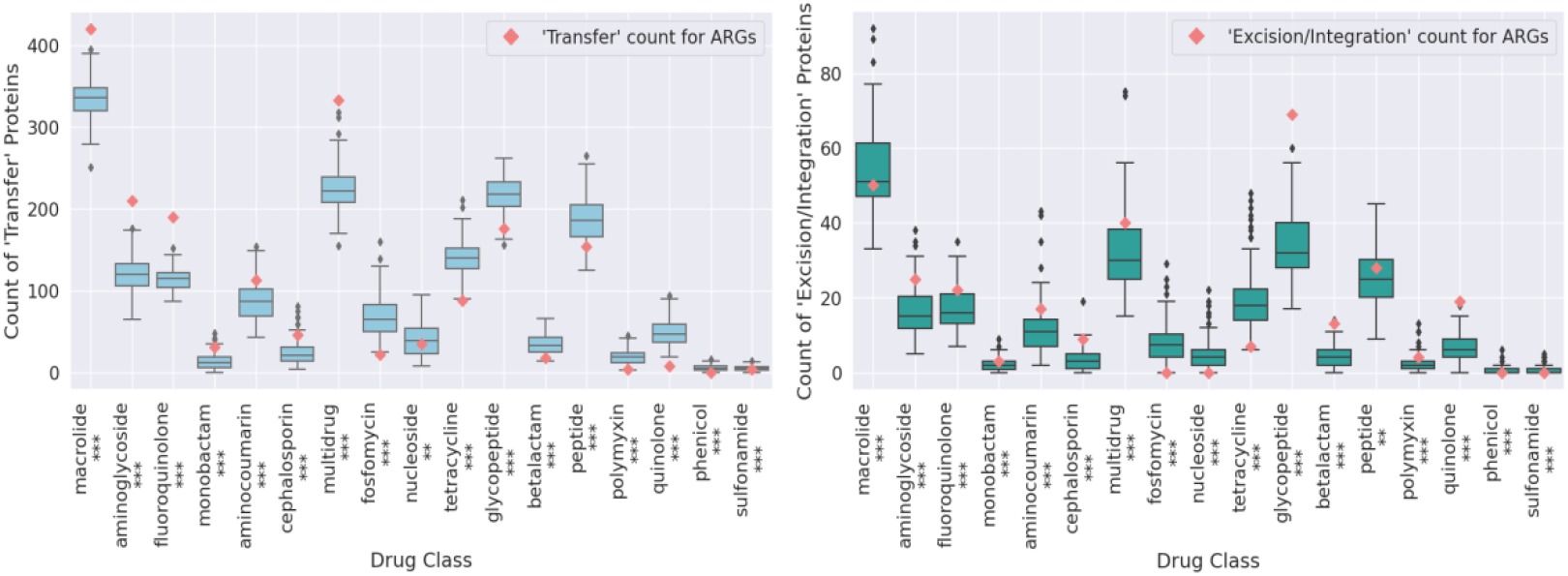
Analysis of the relative mobility of the genes resistant to different drug classes across the ESKAPE pathogens. The distribution of ‘Transfer’ and ‘Excision/Integration’ tags was examined among randomly sampled proteins across various drug classes. The red diamond markers were used to highlight the actual counts of ‘Transfer’ and ‘Excision/Integration’ tags for the ARGs, with asterisks denoting statistical significance (p-value *<* 0.05 based on t-test)

A higher number of excision/integration elements and a lower number of transfer elements were observed in beta-lactam, glycopeptide, peptide, polymyxin, and quinolone drug classes, suggesting that these drug classes may be more prone to gene excision and integration processes, potentially leading to increased resistance. Conversely, we did not observe the opposite pattern in any of the drug classes.

## Discussion and conclusion

PPINs have been used to identify key proteins involved in resistance mechanisms in ESKAPE pathogens and pinpoint essential clusters that highlight significant resistance pathways(34; 35; 36). Additionally, PPINs were employed to shed light on the intricacies of *Staphylococcus aureus* pathogenesis within the context of antibiotic resistance(37). Similarly, the exploration of host-pathogen protein interactomes has proven invaluable, offering a promising avenue for the discovery of novel antibacterial drug targets(38). However, to the best of our knowledge, PPIN and its topological properties have not been employed to develop a machine-learning model for distinguishing ARGs from nonARGs in specific bacterial strains.

Our research builds on our previous work that utilized PPIN analysis in *Escherichia coli* and *Acinetobacter baumannii*(39). We have now developed a pipeline that extends this approach to differentiate ARGs from nonARGs, providing insights into ARG mechanisms and mobility patterns across all ESKAPE pathogens. Additionally, a multiclass classification model was developed as a part of the pipeline to determine the specific drug class to which a resistance gene confers resistance. Importantly, we demonstrated that the ARGs can be classified into different drug classes based solely on their network features, without relying on sequence similarities. Furthermore, through our analysis, we discovered that the neighboring proteins of ARGs exhibited significantly higher mobility compared to nonARGs. This finding suggests a potential link between the mobility of these proteins and the mobility of ARGs. Overall, our study revealed that genes conferring resistance to different drug classes exhibit distinct behaviors within the PPIN with respect to network topology and mobility.

The study also delves into the role of MGEs in spreading antibiotic resistance. The mobility analysis conducted in our study aligns with the initial assumption that ARGs tend to exhibit higher mobility compared to regular nonARGs(40; 41). In contrast, housekeeping genes, being responsible for essential cellular functions, showed less mobility. Moreover, we found that ARG mobility patterns can vary with drug classes; for example, genes resistant to aminoglycosides, aminocoumarins, cephalosporins, monobactams, and multidrugs are more likely to be mobile, correlating with a higher association with MGEs. In contrast, genes resistant to drug classes like sulfonamides, fosfomycin, and phenicol show less mobility, consistent with previous research that reports their chromosomal localization(42; 43). These findings emphasize that the interactions observed in the PPIN provide a rational basis for understanding the mobility of genes resistant to various drug classes.

We conducted our analysis on six highly relevant pathogens known for their mobile forms of antibiotic resistance. In the future, our pipeline can be extended to other virulent and antibiotic-resistant bacterial pathogens. An intriguing aspect of our analysis is the identification of a significant number of proteins falsely predicted as ARGs by our models. This raises questions about their potential roles in resistance mechanisms and whether they could represent novel ARGs, necessitating further investigation. Additionally, our pipeline can be applied to metagenomic data with a preprocessing step required to predict interactions among proteins in different organisms within a metagenomic sample. This approach can enhance existing ARG identification tools and contribute to their robustness. The PPIN pipeline developed in this study can also be applied to identify and study other resistance genes, such as metal resistance genes or biocide resistance genes.

Our pipeline establishes that the network features from the PPIN have some discriminatory power in distinguishing between ARGs and nonARGs. This approach allows for the potential identification of novel ARGs by observing and following the established patterns of well-characterized ARGs within the PPIN. By leveraging PPINs, we gain insights into the mechanisms underlying ARG resistance, and their mobility between bacterial species, and potentially uncover new strategies for combating antibiotic resistance.

## Acknowledgments

This work is supported in part by funds from the National Science Foundation (NSF: #2319521, #2125798, and #2004751).

## References

1. Talebi Bezmin Abadi, A.; Rizvanov, A. A.; Haertlé, T.; Blatt, N. L. World Health Organization Report: Current Crisis of Antibiotic Resistance. BioNanoScience, 4(4):778– 788, 2019.

2. Prestinaci, F.; Pezzotti, P.; Pantosti, A. Antimicrobial Resistance: A Global Multifaceted Phenomenon. Pathogens and Global Health, 7(7):309–318, 2015.

3. Hernando-Amado, S.; Coque, T. M.; Baquero, F.; Martínez, J. L. Defining and Combating Antibiotic Resistance from One Health and Global Health Perspectives. Nature Microbiology, 9(9):1432–1442, 2019.

4. Lushniak, B. D. Antibiotic Resistance: A Public Health Crisis. Public Health Reports, 4(4):314–316, 2014.

5. Centers for Disease Control and Prevention and others. Antibiotic resistance threats in the United States, 2019 US Department of Health and Human Services, Centres for Disease Control.

6. Allen, H. K.; Donato, J.; Wang, H. H.; Cloud-Hansen, K. A.; Davies, J.; Handelsman, J. Call of the Wild: Antibiotic Resistance Genes in Natural Environments. Nature Reviews Microbiology, 4(4):251–259, 2010.

7. Pal, C., Bengtsson-Palme, J., Kristiansson, E. et al. The structure and diversity of human, animal, and environmental resistomes. Microbiome, 4, 54 (2016).

8. Boolchandani, M.; D’Souza, A. W.; Dantas, G. Sequencing-Based Methods and Resources to Study Antimicrobial Resistance. Nature Reviews Genetics, 6(6):356–370, 2019.

9. Berglund, F.; Österlund, T.; Boulund, F.; Marathe, N. P.; Larsson, D. G. J.; Kristiansson, E. Identification and Reconstruction of Novel Antibiotic Resistance Genes from Metagenomes. Microbiome, 1(1):52, 2019.

10. Chen, Q.; Lan, C.; Zhao, L.; Wang, J.; Chen, B.; Chen, Y.-P. P. Recent Advances in Sequence Assembly: Principles and Applications. Briefings in Functional Genomics, 6(6):361–378, 2017.

11. Arango-Argoty, G.; Garner, E.; Pruden, A.; Heath, L. S.; Vikesland, P.; Zhang, L. DeepARG: A Deep Learning Approach for Predicting Antibiotic Resistance Genes from Metagenomic Data. Microbiome, 1(1):23, 2018.

12. Wang, Z.; Li, S.; You, R.; Zhu, S.; Zhou, X. J.; Sun, F. ARG-SHINE: Improve Antibiotic Resistance Class Prediction by Integrating Sequence Homology, Functional Information and Deep Convolutional Neural Network. NAR Genomics and Bioinformatics, 3(3), lqab066, 2021.

13. Ahmed, S.; Emon, M. I.; Moumi, N. A.; Zhang, L. LM-ARG: Identification & Classification of Antibiotic Resistance Genes Leveraging Pre-Trained Protein Language Models. In 2022 IEEE International Conference on Bioinformatics and Biomedicine (BIBM), pages 3782– 3784, 2022.

14. Li, Y.; Xu, Z.; Han, W.; Cao, H.; Umarov, R.; Yan, A.; Fan, M.; Chen, H.; Duarte, C. M.; Li, L.; Ho, P.-L.; Gao, X. H. HMD-ARG: Hierarchical Multi-Task Deep Learning for Annotating Antibiotic Resistance Genes. Microbiome, 1(1):40, 2021.

15. Inda-Díaz, J. S.; Lund, D.; Parras-Moltó, M.; Johnning, A.; Bengtsson-Palme, J.; Kristiansson, E. Latent Antibiotic Resistance Genes Are Abundant, Diverse, and Mobile in Human, Animal, and Environmental Microbiomes. Microbiome, 1(1):44, 2023.

16. Clausen, P. T. L. C.; Zankari, E.; Aarestrup, F. M.; Lund, O. Benchmarking of Methods for Identification of Antimicrobial Resistance Genes in Bacterial Whole Genome Data. Journal of Antimicrobial Chemotherapy, 9(9):2484–2488, 2016.

17. Nielsen, T. K.; Browne, P. D.; Hansen, L. H. Antibiotic Resistance Genes Are Differentially Mobilized According to Resistance Mechanism. GigaScience, 11, giac072, 2022.

18. Slizovskiy, I. B.; Mukherjee, K.; Dean, C. J.; Boucher, C.; Noyes, N. R. Mobilization of Antibiotic Resistance: Are Current Approaches for Colocalizing Resistomes and Mobilomes Useful? Frontiers in Microbiology, 11:1376, 2020.

19. Hanes, R.; Zhang, F.; Huang, Z. Protein Interaction Network Analysis to Investigate Stress Response, Virulence, and Antibiotic Resistance Mechanisms in Listeria Monocytogenes. Microorganisms, 4(4):930, 2023.

20. De Oliveira, D. M. P.; Forde, B. M.; Kidd, T. J.; Harris, P. N. A.; Schembri, M. A.; Beatson, S. A.; Paterson, D. L.; Walker, M. J. Antimicrobial Resistance in ESKAPE Pathogens. Clinical Microbiology Reviews, 3(3):e00181–19, 2020.

21. Denissen, J.; Reyneke, B.; Waso-Reyneke, M.; Havenga, B.; Barnard, T.; Khan, S.; Khan, W. Prevalence of ESKAPE Pathogens in the Environment: Antibiotic Resistance Status, Community-Acquired Infection and Risk to Human Health. International Journal of Hygiene and Environmental Health, 244:114006, 2022.

22. Pendleton, J. N.; Gorman, S. P.; Gilmore, B. F. Clinical Relevance of the ESKAPE Pathogens. Expert Review of Anti Infective Therapy, 3(3):297–308, 2013.

23. von Mering, C.; Jensen, L. J.; Snel, B.; Hooper, S. D.; Krupp, M.; Foglierini, M.; Jouffre, N.; Huynen, M. A.; Bork, P. STRING: Known and Predicted Protein-Protein Associations, Integrated and Transferred across Organisms. Nucleic Acids Research, 33(Database issue):D433–437, 2005.

24. Chen, J.; Chua, H. N.; Hsu, W.; Lee, M.-L.; Ng, S.-K.; Saito, R.; Sung, W.-K.; Wong, L. Increasing Confidence of Protein-Protein Interactomes. Genome Informatics International Conference on Genome Informatics, 2(2):284–297, 2006.

25. Buchfink, B.; Xie, C.; Huson, D. H. Fast and Sensitive Protein Alignment Using DIAMOND. Nature Methods, 1(1):59–60, 2015.

26. Kumar, R. R.; Prasad, S. Metabolic Engineering of Bacteria. Indian Journal of Microbiology, 3(3):403–409, 2011.

27. Petit, C.; Sancar, A. Nucleotide Excision Repair: From E. Coli to Man. Biochimie, 81(1-2):15–25, 1999.

28. Anraku, Y.; Gennis, R. B. The Aerobic Respiratory Chain of Escherichia Coli. Trends in Biochemical Sciences, 12:262–266, 1987.

29. Doncheva, N. T.; Assenov, Y.; Domingues, F. S.; Albrecht, M. Topological Analysis and Interactive Visualization of Biological Networks and Protein Structures. Nature Protocols, 4(4):670–685, 2012.

30. Qi, Y. Random Forest for Bioinformatics. In Ensemble Machine Learning: Methods and Applications; Zhang, C., Ma, Y., Eds.; Springer: New York, NY, 2012; pp 307–323.

31. Brown, C. L.; Mullet, J.; Hindi, F.; Stoll, J. E.; Gupta, S.; Choi, M.; Keenum, I.; Vikesland, P.; Pruden, A.; Zhang, L. mobileOG-Db: A Manually Curated Database of Protein Families Mediating the Life Cycle of Bacterial Mobile Genetic Elements. Applied and Environmental Microbiology, 18(18):e00991–22, 2022.

32. Gilbert, M.; Li, Z.; Wu, X. N.; Rohr, L.; Gombos, S.; Harter, K.; Schulze, W. X. Comparison of Path-Based Centrality Measures in Protein-Protein Interaction Networks Revealed Proteins with Phenotypic Relevance during Adaptation to Changing Nitrogen Environments. Journal of Proteomics, 235:104114, 2021.

33. Li, M.; Zhang, H.; Wang, J.; Pan, Y. A New Essential Protein Discovery Method Based on the Integration of Protein-Protein Interaction and Gene Expression Data. BMC Systems Biology, 1(1):15, 2012.

34. Priyamvada, P.; Debroy, R.; Anbarasu, A.; Ramaiah, S. A Comprehensive Review on Genomics, Systems Biology and Structural Biology Approaches for Combating Antimicrobial Resistance in ESKAPE Pathogens: Computational Tools and Recent Advancements. World Journal of Microbiology and Biotechnology, 9(9):153, 2022.

35. Miryala, S. K.; Ramaiah, S. Exploring the Multi-Drug Resistance in Escherichia Coli O157:H7 by Gene Interaction Network: A Systems Biology Approach. Genomics, 4(4):958–965, 2019.

36. Debroy, R.; Miryala, S. K.; Naha, A.; Anbarasu, A.; Ramaiah, S. Gene Interaction Network Studies to Decipher the Multi-Drug Resistance Mechanism in Salmonella Enterica Serovar Typhi CT18 Reveal Potential Drug Targets. Microbial Pathogenesis, 142:104096, 2020.

37. Otarigho, B.; Falade, M. O. Analysis of Antibiotics Resistant Genes in Different Strains of Staphylococcus Aureus. Bioinformation, 3(3):113–122, 2018.

38. Zoraghi, R.; Reiner, N. E. Protein Interaction Networks as Starting Points to Identify Novel Antimicrobial Drug Targets. Current Opinion in Microbiology, 5(5):566–572, 2013.

39. Moumi, N. A.; Brown, C. L.; Vikesland, P. J.; Pruden, A.; Zhang, L. Protein-Protein Interaction Network Analysis Reveals Distinct Patterns of Antibiotic Resistance Genes. In 2022 IEEE International Conference on Bioinformatics and Biomedicine (BIBM), pages 73–76, 2022.

40. Stokes, H. W.; Gillings, M. R. Gene Flow, Mobile Genetic Elements and the Recruitment of Antibiotic Resistance Genes into Gram-Negative Pathogens. FEMS Microbiology Reviews, 5(5):790–819, 2011.

41. Partridge, S. R.; Kwong, S. M.; Firth, N.; Jensen, S. O. Mobile Genetic Elements Associated with Antimicrobial Resistance. Clinical Microbiology Reviews, 4(4):e00088–17, 2018.

42. Rizzo, L.; Manaia, C.; Merlin, C.; Schwartz, T.; Dagot, C.; Ploy, M. C.; Michael, I.; Fatta-Kassinos, D. Urban Wastewater Treatment Plants as Hotspots for Antibiotic Resistant Bacteria and Genes Spread into the Environment: A Review. Science of the Total Environment, 447:345–360, 2013.

43. Li, W.; Mao, F.; Ng, C.; Jong, M. C.; Goh, S. G.; Charles, F. R.; Ng, O. T.; Marimuthu, K.; He, Y.; Gin, K. Y.-H. Population-Based Variations of a Core Resistome Revealed by Urban Sewage Metagenome Surveillance. Environmental International, 163:107185, 2022.

