## Supplementary Material for "Resistance genes are distinct in protein-protein interaction networks according to drug class and gene mobility"

Supplementary Fig. 1

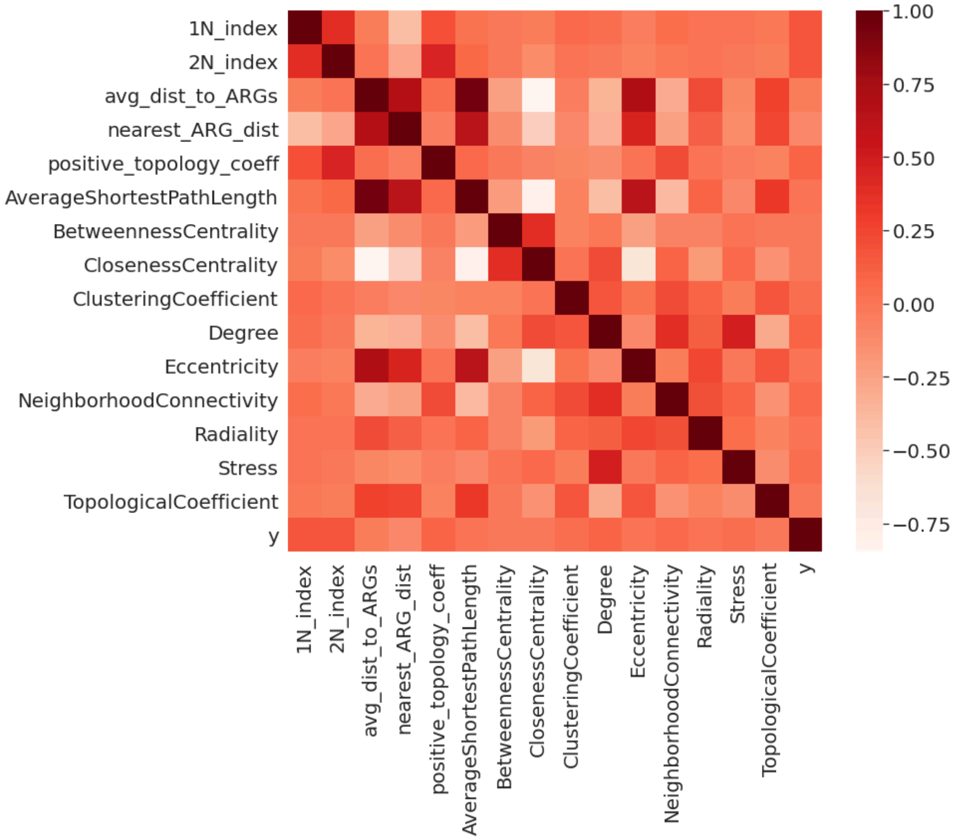


Supplementary Fig. 1: Feature correlation for the 15 features across the combined set of proteins from all six ESKAPE pathogens.

Supplementary Fig. 2

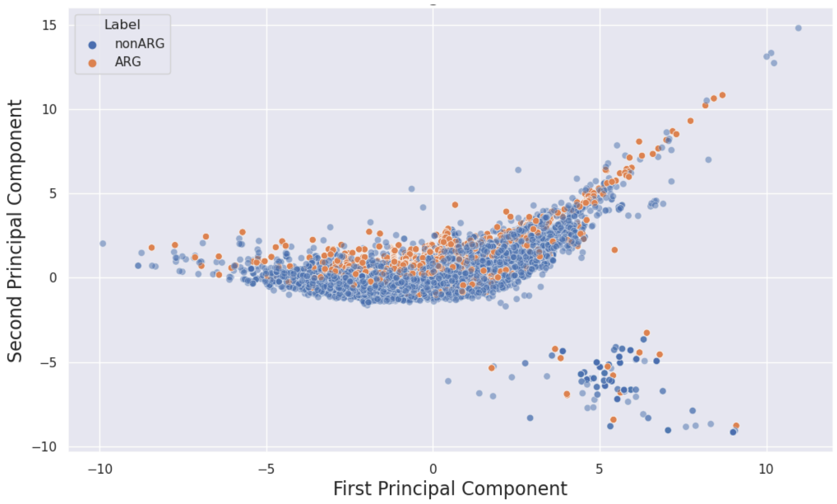


Supplementary Fig. 2: Principal components analysis (PCA) plot for the combined set of proteins from all six ESKAPE pathogens. Orange dots denote the ARGs, and blue dots represent the nonARGs.

Supplementary Fig. 3

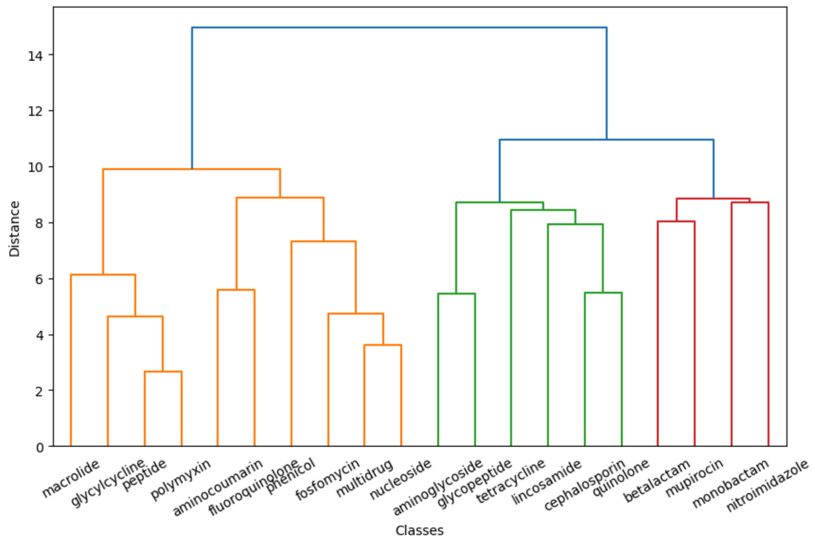


Supplementary Fig. 3: Dendrogram representing  single linkage hierarchical clustering of the top 20 most frequent drug classes based on the feature values from the PPIN

Supplementary Fig. 4

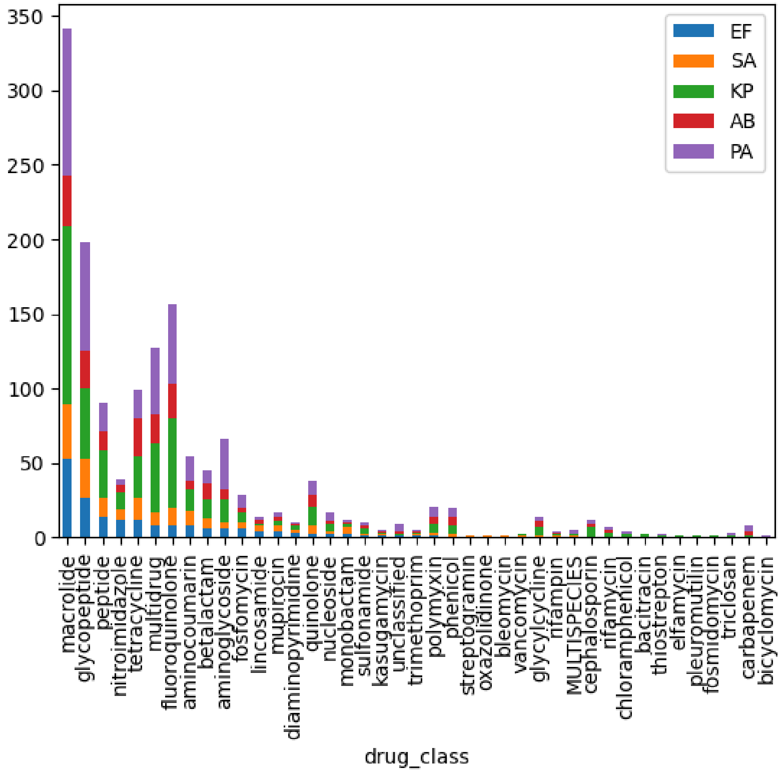


Supplementary Fig. 4: Bar plot depicting the actual counts of resistance genes, displayed on the x-axis, for each drug class represented on the y-axis. The plot features five distinct colors to represent five pathogens: (Blue: *Enterococcus faecium*, orange: *Staphylococcus aureus*, green: *Klebsiella pneumoniae*, Red: *Acinetobacter baumannii*, purple: *Pseudomonas aeruginosa*)

Supplementary Table 1

| |
|---|
| aerobic respiration [GO:0009060] |
| carbon catabolite repression of transcription [GO:0045013] |
| transcription-coupled nucleotide-excision repair [GO:0006283] |
| transcription-coupled nucleotide-excision repair, DNA damage recognition [GO:0000716] |
| nucleotide-excision repair, DNA damage recognition [GO:0000715] |
| nucleotide-excision repair, preincision complex assembly [GO:0006294] |
| base-excision repair, DNA ligation [GO:0006288] |
| nucleotide-excision repair [GO:0006289] |
| base-excision repair, AP site formation via deaminated base removal [GO:0097510] |
| nucleotide-excision repair, DNA incision [GO:0033683] |
| base-excision repair [GO:0006284] |
| nucleotide-excision repair, DNA duplex unwinding [GO:0000717] |
| base-excision repair, AP site formation [GO:0006285] |
| canonical glycolysis [GO:0061621] |
| acyl-CoA metabolic process [GO:0006637] |
| one-carbon metabolic process [GO:0006730] |
| lipoprotein metabolic process [GO:0042157] |
| regulation of RNA metabolic process [GO:0051252] |
| hypoxanthine metabolic process [GO:0046100] |
| amine metabolic process [GO:0009308] |
| inositol metabolic process [GO:0006020] |
| UDP-glucose metabolic process [GO:0006011] |
| ribose phosphate metabolic process [GO:0019693] |
| sulfur amino acid metabolic process [GO:0000096] |
| 3'-phosphoadenosine 5'-phosphosulfate metabolic process [GO:0050427] |
| regulation of glutamine family amino acid metabolic process [GO:0000820] |
| dodecyl sulfate metabolic process [GO:0018909] |
| fructose 1,6-bisphosphate metabolic process [GO:0030388] |
| nitrate metabolic process [GO:0042126] |
| glycerol to glycerone phosphate metabolic process [GO:0061610] |
| glutathione metabolic process [GO:0006749] |
| glucose 6-phosphate metabolic process [GO:0051156] |

GDP-mannose metabolic process [GO:0019673]

carbohydrate derivative metabolic process [GO:1901135]

regulation of cellular organohalogen metabolic process [GO:0090347]

biotin metabolic process [GO:0006768]

succinate metabolic process [GO:0006105]

aminophosphonate metabolic process [GO:0033051]

N-acetylneuraminate metabolic process [GO:0006054]

siderophore metabolic process [GO:0009237]

nucleobase-containing compound metabolic process [GO:0006139]

dihydrofolate metabolic process [GO:0046452]

folic acid-containing compound metabolic process [GO:0006760]

glutamine metabolic process [GO:0006541]

L-lyxose metabolic process [GO:0019324]

IMP metabolic process [GO:0046040]

galactitol metabolic process [GO:0019402]

shikimate metabolic process [GO:0019632]

D-amino acid metabolic process [GO:0046416]

peptidoglycan metabolic process [GO:0000270]

nitroglycerin metabolic process [GO:0018937]

nucleotide metabolic process [GO:0009117]

hydrogen peroxide metabolic process [GO:0042743]

S-methylmethionine metabolic process [GO:0033477]

regulation of nitrogen compound metabolic process [GO:0051171]

cellular aromatic compound metabolic process [GO:0006725]

positive regulation of secondary metabolite biosynthetic process [GO:1900378]

regulation of tryptophan metabolic process [GO:0090357]

positive regulation of cellular carbohydrate metabolic process [GO:0010676]

nucleoside phosphate metabolic process [GO:0006753]

L-ascorbic acid metabolic process [GO:0019852]

succinyl-CoA metabolic process [GO:0006104]

thiamine metabolic process [GO:0006772]

fucose metabolic process [GO:0006004]

amino sugar metabolic process [GO:0006040]

coenzyme A metabolic process [GO:0015936]

| |
|---|
| purine nucleotide metabolic process [GO:0006163] |
| D-xylose metabolic process [GO:0042732] |
| sulfur compound metabolic process [GO:0006790] |
| heme metabolic process [GO:0042168] |
| glutamate metabolic process [GO:0006536] |
| glycerol-3-phosphate metabolic process [GO:0006072] |
| glucose metabolic process [GO:0006006] |
| malate metabolic process [GO:0006108] |
| positive regulation of carbohydrate metabolic process [GO:0045913] |
| alpha-amino acid metabolic process [GO:1901605] |
| phosphatidylglycerol metabolic process [GO:0046471] |
| lipoate metabolic process [GO:0009106] |
| thymidine metabolic process [GO:0046104] |
| secondary metabolite biosynthetic process [GO:0044550] |
| pyrimidine nucleobase metabolic process [GO:0006206] |
| UDP-N-acetylglucosamine metabolic process [GO:0006047] |
| trehalose metabolism in response to cold stress [GO:0070415] |
| pyrimidine nucleotide metabolic process [GO:0006220] |
| fatty acid metabolic process [GO:0006631] |
| fumarate metabolic process [GO:0006106] |
| nucleobase-containing small molecule metabolic process [GO:0055086] |
| rhamnose metabolic process [GO:0019299] |
| peptide metabolic process [GO:0006518] |
| carboxylic acid metabolic process [GO:0019752] |
| acetate metabolic process [GO:0006083] |
| NADP metabolic process [GO:0006739] |
| N-acetylglucosamine metabolic process [GO:0006044] |
| diacetylchitobiose metabolic process [GO:0052778] |
| carbohydrate metabolic process [GO:0005975] |
| D-gluconate metabolic process [GO:0019521] |
| cellular carbohydrate metabolic process [GO:0044262] |
| selenocysteine metabolic process [GO:0016259] |
| acetyl-CoA metabolic process [GO:0006084] |
| argininosuccinate metabolic process [GO:0000053] |

galactarate metabolic process [GO:0019580]

guanosine tetraphosphate metabolic process [GO:0015969]

generation of precursor metabolites and energy [GO:0006091]

NADH metabolic process [GO:0006734]

negative regulation of cellular carbohydrate metabolic process [GO:0010677]

aspartate metabolic process [GO:0006531]

nitrogen compound metabolic process [GO:0006807]

colanic acid metabolic process [GO:0046377]

organic phosphonate metabolic process [GO:0019634]

superoxide metabolic process [GO:0006801]

pyrimidine nucleoside metabolic process [GO:0006213]

L-fucose metabolic process [GO:0042354]

nucleoside monophosphate metabolic process [GO:0009123]

glycerol metabolic process [GO:0006071]

fructose metabolic process [GO:0006000]

metabolic process [GO:0008152]

toxic metabolite repair [GO:0110052]

propionate metabolic process, methylcitrate cycle [GO:0019679]

threonine metabolic process [GO:0006566]

negative regulation of carbohydrate metabolic process [GO:0045912]

tetrahydrofolate metabolic process [GO:0046653]

phosphatidylcholine metabolic process [GO:0046470]

purine nucleobase metabolic process [GO:0006144]

phosphorus metabolic process [GO:0006793]

citrate metabolic process [GO:0006101]

lipid metabolic process [GO:0006629]

D-glucarate metabolic process [GO:0042836]

trehalose metabolic process [GO:0005991]

D-serine metabolic process [GO:0070178]

pyridoxal metabolic process [GO:0042817]

nucleoside diphosphate metabolic process [GO:0009132]

fructose 6-phosphate metabolic process [GO:0006002]

lipopolysaccharide metabolic process [GO:0008653]

N-acetylmannosamine metabolic process [GO:0006051]

tartrate metabolic process [GO:1901275]

2-oxoglutarate metabolic process [GO:0006103]

regulation of fatty acid metabolic process [GO:0019217]

xenobiotic metabolic process [GO:0006805]

negative regulation of phosphate metabolic process [GO:0045936]

D-ribose metabolic process [GO:0006014]

riboflavin metabolic process [GO:0006771]

ferredoxin metabolic process [GO:0006124]

oxaloacetate metabolic process [GO:0006107]

cytosine metabolic process [GO:0019858]

positive regulation of tryptophan metabolic process [GO:0090358]

NAD metabolic process [GO:0019674]

primary metabolic process [GO:0044238]

protoporphyrinogen IX metabolic process [GO:0046501]

cellular metabolic compound salvage [GO:0043094]

carnitine metabolic process [GO:0009437]

propanediol metabolic process [GO:0051143]

regulation of secondary metabolite biosynthetic process [GO:1900376]

glyoxylate metabolic process [GO:0046487]

mannitol metabolic process [GO:0019594]

short-chain fatty acid metabolic process [GO:0046459]

formaldehyde metabolic process [GO:0046292]

dimethyl sulfoxide metabolic process [GO:0018907]

metabolite repair [GO:0110051]

glycerophospholipid metabolic process [GO:0006650]

pyruvate metabolic process [GO:0006090]

phosphate-containing compound metabolic process [GO:0006796]

galactose metabolic process [GO:0006012]

purine nucleoside metabolic process [GO:0042278]

ribonucleoside diphosphate metabolic process [GO:0009185]

cellular amino acid metabolic process [GO:0006520]

mannose metabolic process [GO:0006013]

cellular lipid metabolic process [GO:0044255]

nicotinamide nucleotide metabolic process [GO:0046496]

| |
|---|
| homocysteine metabolic process [GO:0050667] |
| folic acid metabolic process [GO:0046655] |
| tRNA metabolic process [GO:0006399] |
| chorismate metabolic process [GO:0046417] |
| L-serine metabolic process [GO:0006563] |
| guanine metabolic process [GO:0046098] |
| nucleotide-sugar metabolic process [GO:0009225] |
| nucleoside metabolic process [GO:0009116] |
| arginine metabolic process [GO:0006525] |
| spermidine metabolic process [GO:0008216] |
| cellular biogenic amine metabolic process [GO:0006576] |
| DNA metabolic process [GO:0006259] |
| positive regulation of isoprenoid metabolic process [GO:0045828] |
| pentose-phosphate shunt [GO:0006098] |
| pentose-phosphate shunt, oxidative branch [GO:0009051] |
| pentose-phosphate shunt, non-oxidative branch [GO:0009052] |

Supplementary Table 1: GO terms for the housekeeping genes
